# Rab35 drives epidermal wound closure by promoting lamellipodial protrusions and membrane vesicle transport

**DOI:** 10.64898/2026.10.08.757533

**Authors:** Meike Schneider, Sven Bogdan

## Abstract

In response to tissue damage, the epidermal barrier must be restored immediately to prevent excessive blood loss and infections. How cells coordinate actin-driven protrusions and membrane transport during reepithelialization is poorly understood. Here, we identified Rab35 as an important driver of epidermal wound closure in *Drosophila*. Endogenous Rab35 positive vesicles are directed to the wound edge and in protruding lamellipodia. Suppression of Rab35 function results in striking lamellipodia defects and an accumulation of PIP2-positive vesicles. Depletion of Fascin, a known Rab35 effector phenocopies *rab*35 knockdown. Upstream of Rab35, we identified CG18659 as potential Rab35-GEF protein, which physically interacts and colocalizes with Rab35. Suppression of Rab35 GEF CG18659 or overexpression of Rab35 GAP Whacked results in strong wound healing defects. We propose model in which Rab35 drives epidermal wound closure by promoting Fascin-stabilized actin-driven protrusions and vesicle transport required for cell membrane expansion and local PIP2 delivery at the wound site.

**Significance statement:** To restore the epidermal barrier and protect the body, the skin immediately triggers a highly coordinated wound healing process. How cells dynamically coordinate actin-driven protrusions and membrane transport during epidermal wound closure is poorly understood. Here, we identified the conserved Rab35 as a central driver in epidermal wound healing coordinating both structural actin remodeling and localized phosphoinositide signaling, required for efficient tissue repair. Our results give a new perspective on RabGTPases with dual functions such as Rab35 synchronize actin-driven protrusions and membrane transport during epidermal wound closure.

## Introduction

Epithelia form highly selective barriers lining most of our organs. Upon injury, epithelial tissues must seal rapidly and remodel the wounded area in order to prevent infections and to maintain tissue function ^1^. Thus, wound healing is an essential physiological process that has been studied extensively in different model systems ^2^. Despite morphological differences found in distinct tissues and developmental stages the process of wound healing comprises sequential steps that are controlled by conserved molecular and cellular mechanisms across species ^3^ ^4^. During wound healing, epithelial cells migrate collectively as a coherent sheet to heal the wound that involves a rapid reorganization of the actin cytoskeleton at the wound edge ^5–11^. Collective migration of epithelial cells depends not only on the precise coordination of actin-driven protrusion dynamics, but also on vesicle trafficking, which continuously cycle membrane lipids and cell surface receptors to the leading edge of the migrating cohort ^12,13^. However, the role of membrane vesicle transport in epithelial wound healing are not well understood *in vivo*, and the molecular mechanisms and drivers remain largely unclear. The Ras-associated binding (Rab) family of GTPases are the primary regulators of the vesicular trafficking pathways that are responsible for transporting the vast array of cellular cargo across ^14–17^. As shown for other monomeric small GTPases, Rab proteins bind GTP and hydrolyze it, switching between inactive and active states through the binding of GDP or GTP, a process regulated by GAPs or GEFs ^18^. The Rab family comprises approximately 70 members in humans differentially regulating intracellular membrane traffic in cell polarity, cytokinesis, growth factor signaling, and cell migration^15^. Mutations in *rab* genes can cause a wide range of genetic diseases, ranging from neurodegenerative diseases to immune deficiency syndromes and to a variety of cancers ^19^. A role of distinct Rab proteins, however, has so far been described only *in vitro* wound healing assays ^20,21^. Using the *Drosophila* larval epidermis as an *in vivo* wound model ^9^, we performed an RNAi knockdown candidate screen for 32 of the 33 *rab* genes encoded in the in *Drosophila* genome. Among others, we identified the conserved Rab35 GTPase as the strongest candidate required for proper epidermal wound closure. Upon wounding, endogenous Rab35 localizes to endosomal vesicles, which are transported to the wound edge and into newly formed protruding lamellipodia. Consequently, suppression of Rab35 function results in an accumulation of phosphatidylinositol 4,5-bisphosphate (PIP2)-positive vesicles as well as in prominent lamellipodia defects. Upstream of Rab35, we identified the DENN domain containing GEF protein CG18659, which physically interacts with Rab35. Suppression of CG18659 and the actin-bundling protein Fascin, a known Rab35 downstream effector phenocopies *rab*35 depletion. Thus, we propose a model in which Rab35 drives epidermal wound closure by coordinating actin-driven lamellipodial protrusions and the polarized transport of PIP2-containing vesicles required for cell membrane expansion and gap closure at the wound edge.

## Results

### Rab35 acts through Fascin to regulate lamellopodial protrusions in wound closure

The *Drosophila* larval epidermis served as a powerful genetic model system to identify conserved genes that are required for normal wound healing ^2,9,22^. The postmitotic so-called larval epidermal cells (LECs) are up to 100 µm in diameter and persist until early pupal metamorphosis (Figure 1a, b), make them an attractive model for studying cellular dynamics in single-cell wounding experiments at high resolution ^22,23^. The main actin-driven structures that contribute to wound closure are protruding Rac-WAVE-Arp2/3 dependent lamellipodia and actomyosin driven contractile cables which form in different spatially constrained regions of the adjacent cells (Figure 1b), found both in invertebrates and vertebrates ^9,24^. Live-cell microscopy of laser-induced epidermal wounds also reveals numerous dynamic vesicular structures that may contribute to efficient epidermal wound closure (supplementary movie 1). A critical contribution of intracellular membrane trafficking in epithelial wound healing has been proposed but the molecular mechanisms and drivers remain largely unknown. To better understand the role of membrane trafficking in wound healing, we performed an RNAi-based candidate approach targeting 32 of the 33 *Drosophila* Rab GTPases, which are considered to be key organizers in vesicular trafficking ^25^. Two different RNAi lines were tested for each *rab* gene (Supplementary table 1). To this aim, transgenic RNAi lines were co-expressed with a Lifeact-EGFP transgene recombined with the epidermis-specific A58-Gal4 driver ^26^. Lifeact-GFP was used as a reporter to visualize changes in actin dynamics and wound closure efficiency (Figure 1c). Knockdown of Rab GTPases resulted in different wound closure defects which were classified as a strong, weak or no phenotype (Supplementary table 2). Among them, RNAi-mediated depletion of *rab35* using two independent RNAi lines resulted in striking changes in F-actin dynamics and in strong wound closure defects (Figure 1d, e). In both cases, we observed an overall reduction in cortical F-actin (Figure 1d, e). Moreover, we observed an accumulation of numerous F-actin coated vesicular structures and the formation of dynamic actin comet tails racing through epidermal cells (Figure 1d, e, yellow arrow heads; supplementary movies M2 and M3). Compared to wild type, cells at the wound margin depleted for *rab35* showed reduced lamellipodia size and a strong delay in wound healing within the first 30 minutes (Figure 1d, e; quantification in Figure 1g and h). Upon overexpression of a dominant-negative Rab35DN locking it in an inactive, GDP-bound state ^27^, we observed similar wound closure defects, with more cryptic lamellipodia, abnormal vesicle accumulations and intense actin comet tail formation (Figure 1f; quantification in Figure 1g and h; supplementary movie M4).

**Figure 1:**
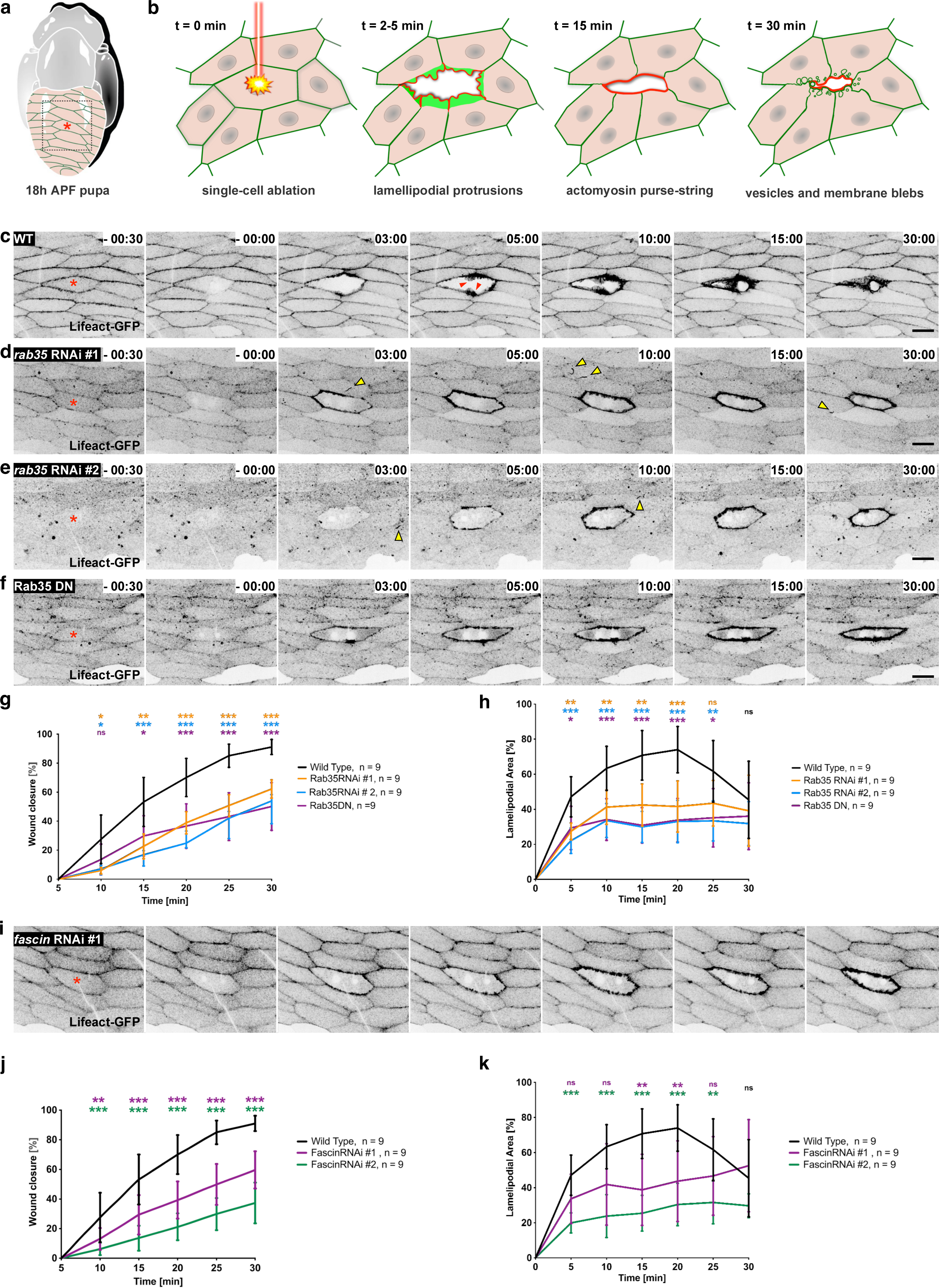
Rab35 acts through Fascin to regulate lamellopodial protrusions in wound closure. **(a)** Schematic of a wild type pupa, 18h APF, expressing a green-fluorescent Lifeact-GFP reporter in the abdominal epidermis. The red asterisk marks single-cell laser ablation. **(b)** Zoom in from (a). Schematic of an *in vivo* model for laser-induced wounding. Single-cell laser ablation begins at t = 0 min. After two minutes (t = 2 min), F-actin accumulates into lamellipodial protrusions (green) within cells at the wound margin. These lamellipodia reach their maximum extend between 5 and 10 minutes (t = 5 min) after wounding. At around 15 minutes (t = 15 min) an actomyosin purse string (red) forms at the leading edge of the wound zipping opposite wound edges together to drive wound closure. Later on (t = 30 min) vesicles and membrane blebs form around the wound margin. (**c-f**) Single frames of time-lapse confocal spinning disc movies. A Lifeact-EGFP transgene was expressed in the abdominal epidermis, 18h APF, under control of the A58-Gal4 driver. Frames were taken at indicated time points. Single-cell laser ablation (red asterisk) starts at t = 0 min. The genotypes are indicated: **(c)** In the wild type (WT), a broad lamellipodium is formed (n = 9) and the wound has been closed after 30 min, whereas knockdown of **(d, e)** Rab35 (n = 9 each) and **(f)** dominant-negative Rab35DN (n = 9) resulted in cryptic lamellipodia. Scale bar: 20 µm. **(g)** Quantification of wound closure in wild type (n = 9) and after knockdown of Rab35 using two independent RNAi lines (n = 9 each) as well as dominant-negative Rab35DN (n = 9) over the time for 30 min. **(h)** Quantification of the lamellipodial area for wild type (n = 9), after knockdown of Rab35 using RNAi (n = 9) as well as for dominant-negative Rab35DN (n = 9). The lamellipodial area was measured every 5 min. **(i)** Frames of a single-cell wound healing time-lapse movie. Frames were taken at indicated time points. *fascin* was knocked down using RNAi. A58-Gal4-dependent expression of Lifeact-EGFP in an 18h APF abdominal epidermis. Single-cell laser ablation (red asterisk) started at *t = 0 min*. Scale bar: 20 µm. **(j)** Quantification of wound closure within 30 min in wild type (n = 9) and after depletion of Fascin using two different RNAi lines (n = 9 each). **(k)** Quantification of the lamellipodial area for wild type (n = 9) and *fascin* RNAi (n = 9 each). Note that wild type controls are the same as in Fig. 1g and h as the data belong to the same dataset. The lamellipodium was measured every 5 minutes. At least three independent experiments for each genotype in c-f were performed. The statistical significance in g, h, j and k was evaluated using the two-way ANOVA analysis with Geisser-Greenhouse correction and Dunnett‘s multiple comparisons test. The following p-values were obtained: *p = 0.12* (ns), *p = 0.033* (*), *p = 0.002* (**) and *p < 0.001* (***).

Since active Rab35 regulates actin bundling by recruiting the actin-bundling protein Fascin as an effector driving lamellipodial protrusions ^28^, we next analyzed a possible role of Fascin downstream of Rab35 in epidermal wound closure. Depletion of Fascin by using two independent RNAi transgenes resulted indeed in prominent defects in lamellipodia formation and a significant delay in wound healing which resembles Rab35 depletion (Figure 1i; quantification in Figure 1j and k; supplementary movie M5). Different from *rab35* depleted epidermal cells, however, no vesicle accumulations and intense actin comet tail formation were observed Figure 1i; supplementary movie M5). This suggests that Rab35 acts through Fascin to regulate actin-driven lamellopodial protrusions in wound closure but that this interaction plays no role in vesicle transport or vesicle movement by actin comet tails.

### Rab35 is regulated by the guanine-nucleotide exchange factor (GEF) CG18659 and the GTPase-activating protein (GAP) Whacked in wound healing

Rab proteins are activated by guanine-nucleotide exchange factors (GEFs) that mediate the switch from GDP to GTP and are inactivated through GTPase-activating proteins (GAPs) that promote the intrinsic GTPase activity ^25,29^. Three potential Rab35 GEFs (CG18659, Stratum/CG7787, and Sbf) were biochemically identified using affinity chromatography in *Drosophila* ^30^ and two distinct GAP proteins, Skywalker (Sky) and Whacked (Wkd) that differentially stimulate the intrinsic GTPase activity of Rab35 in neurons and epithelial cells, respectively ^31,32^. RNAi-mediated depletion of CG18659 phenocopied indeed *rab35* knockdown (Figure 2a; quantification in Figure 2d). Cells depleted for CG18659 also formed cryptic lamellipodia at wound margin and actin comet tails similar to those observed in a *rab35* knockdown, albeit somewhat less pronounced (Figure 2a, d; supplementary movie M6). Suppression of Whacked function did not impair wound healing; on the contrary, the cells formed very large lamellipodia, and wound closure occurred very efficiently (Figure 2b, d; supplementary movie M7). By contrast, overexpression of wild type Whacked that inactivates Rab35 resulted in strong lamellipodia and wound closure defects (Figure 2c, d; supplementary movie M8). Thus, these data suggest that Rab35 function is regulated upstream by the potential GEF CG18659 and the known GAP Whacked in epidermal wound healing. CG18659 belongs to the highly conserved tripartite DENN (differentially expressed in normal and neoplastic cells) domain–containing Rab-specific guanine nucleotide exchange factors ^33,34^. The DENN domain consists of three subdomains: uDENN (upstream DENN), the core DENN domain, and dDENN, which are structurally distinct but functionally act together enzymatically as a Rab-specific guanine nucleotide exchange factor (Figure 1e). The predicted AlphaFold 3 structure of *Drosophila* CG18659 resembled the known crystal structure of the human orthologues DENND1A-C ^35^. AlphaFold 3 also predicted that CG18659 is able to directly bind Rab35 with high confidence (ipTM=0.84; pTM=0.73). Co-immunoprecipitations with extracts of *Drosophila* S2R+ cells co-transfected with GFP-tagged Rab35 and with a Myc-tagged CG18659 further confirmed a physical interaction (Figure 2g). In cultured S2R+ cells that had spread out on surfaces coated with concanavalin A (Con A), transfected Rab35 partially co-localized with F-actin-rich structures (Figure 2h, whereas a Myc-tagged CG18659 protein was largely found at the plasma membrane (Figure 2i). Co-transfection of both constructs resulted in a partial colocalization of Rab35 and CG18659 at the protruding leading edge of S2R+ cells (Figure 1j). Thus, Rab35 and its potential GEF, CG18659, interact with each other and determine their cellular localization.

**Figure 2:**
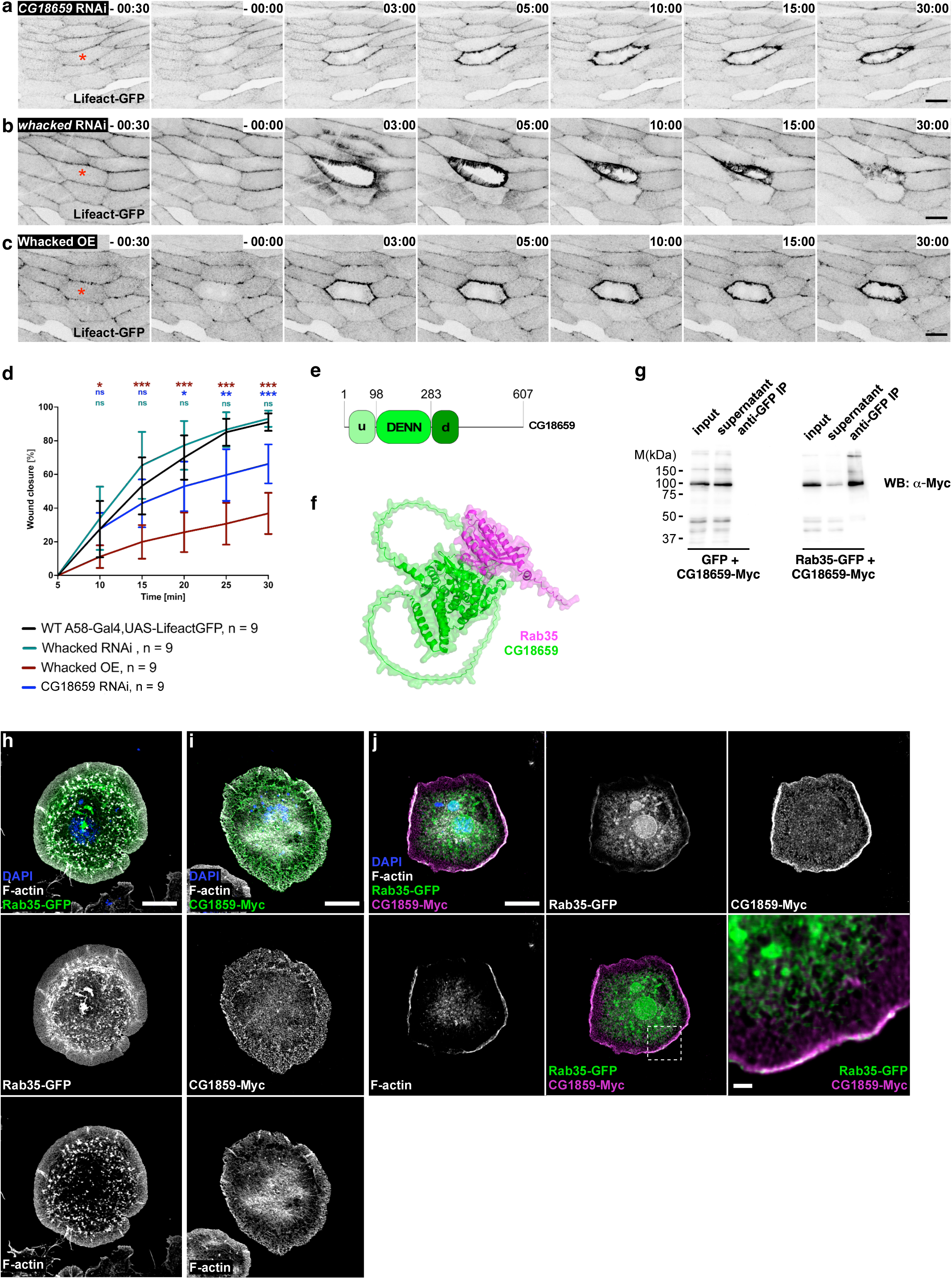
Rab35 is regulated by the guanine-nucleotide exchange factor (GEF) CG18659 and the GTPase-activating protein (GAP) Whacked in wound healing (a-c) Single frames of spinning disc microscope videos from single-cell laser ablation experiments in the 18h APF old abdominal epidermis after knockdown of CG18659 **(a)**, after knockdown of *whacked* (b) or after overexpression of Whacked **(c)**. The epidermis expressed Lifeact-EGFP under the control of A58-Gal4 driver. Images were taken at indicated time points. Ablation of a single cell (red asterisk) starts at t = 0 min. Scale bar: 20 µm. **(d)** Quantification of wound closure over the time of 30 minutes in wild type (WT, n = 9) as well as when CG18659 (GEF) or whacked were knocked down using RNAi or whacked was overexpressed (OE). Note that wild type controls are the same as in Fig. 1g as the data belong to the same dataset. The statistical significance was tested using the two-way ANOVA analysis with Geisser-Greenhouse correction and Dunnett‘s multiple comparisons test. The following p-values were obtained: p = 0.12 (ns), p = 0.033 (*), p = 0.002 (**) and p < 0.001 **(e)** Schematic domain structure of CG18659 containing the tripartite DENN domain important for GEF activity. **(e)** AlphaFold3-modelled protein structure of CG18659 (green) bound to GTP-Rab35 (magenta). The confidence metrics are: ipTM = 0.84 and pTM = 0.73. **(g)** Western Blot from a co-immunoprecipitation of Rab35-GFP with CG18659-myc, stained against anti-myc. GFP was pulled down. The input fraction was 2 % of the cell lysate. CG18659-myc binds to Rab35-GFP. **(h-j)** Confocal images of transfected, fixed and stained *Drosophila* S2R+ cells. All cells were stained with DAPI (nuclei) and Phalloidin (F-actin). **(h)** S2R+ cells transfected with Rab35-GFP. **(i)** S2R+ cells transfected with CG18659-myc. **(j)** S2R+ cell co-transfected with Rab35-GFP and CG18659-myc which was detected with an anti-myc antibody. At least three independent experiments for each genotype in a-c were performed. Co-IP and S2R+ cell transfection were performed in triplicates. Scale bar represents 10 µm and 2 µm in the zoom shown in **(j)**.

### Endogenous Rab35 positive vesicles are directed to the wound edge and in protruding lamellipodia during epidermal wound closure

To visualize Rab35 localization in epidermal cells upon wounding *in vivo* we made use of an endogenously marked YFP-Rab35 transgene previously generated by the Brankatschk lab ^36^. In unwounded LECs, endogenous Rab35 is distributed in puncta and along cell-cell interfaces on the apical surface (Figure 3a; supplementary movie M9). Following laser-induced wound wounding, Rab35 accumulates extensively in numerous vesicles that appear near the wound margin (Figure 3a; supplementary movie M9). During wound closure, an increasing number of Rab35-positive vesicles fuse with the apical membrane, which becomes increasingly visible over time as a thin line (Figure 3a; supplementary movie M9). Starting from the apical wound edge, Rab35 also accumulated increasingly into lamellipodial protrusions (Figure 3a; supplementary movie M9). Tracking of individual Rab35-positive vesicles confirmed a polarized movement predominantly toward the wound margin (Figure 3b, c).

**Figure 3:**
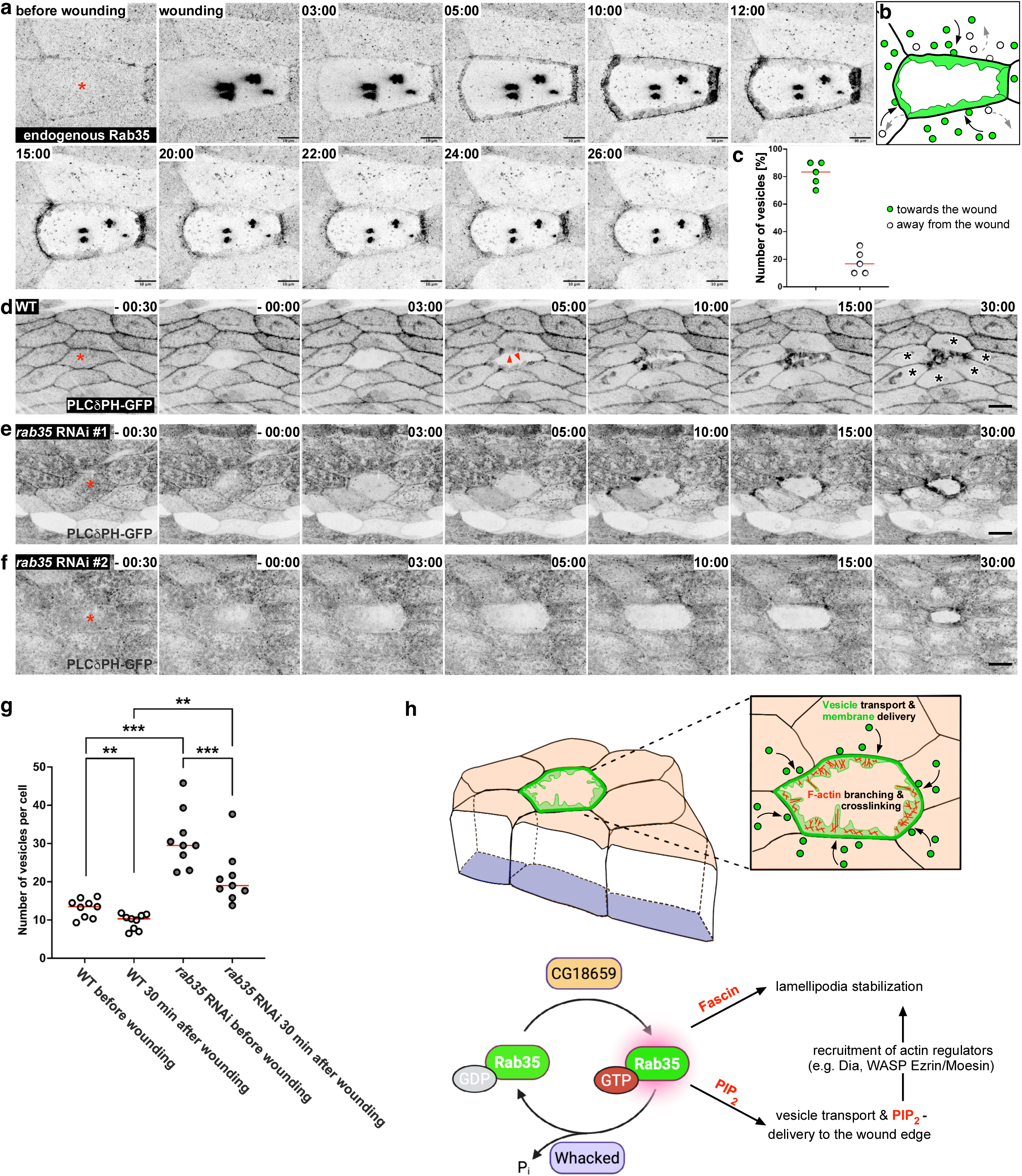
Suppression of Rab35 function affects vesicular transport of PIP2. **(a)** Frames of an airy scan confocal microscope time-lapse video from single-cell wounding experiments in an 18h APF old abdominal epidermis, expressing endogenous Rab35 (n = 5). Images were taken at indicated time points. Ablation of a single cell (red asterisk) starts at t = 0 min. Scale bar represents 10 µm. **(b)** Schematic of vesicle movement from (a). Green vesicles move towards the wound margin whereas white vesicles move away from the wound margin. **(c)** Quantification of vesicle movement from n =5 movies. **(d-f)** Single frames of spinning disc microscope time lapse videos from single-cell laser ablation experiments in the 18h APF old abdominal epidermis, expressing PLCδPH-GFP under control of A58-Gal4 in wild type (WT, n = 9) or when Rab35 was knocked down using RNAi (n = 9). Images were taken at indicated time points. Ablation of a single cell (red asterisk) starts at t = 0 min. Scale bar represents 20 µm. **(g)** Quantification of PIP2-positive vesicles before (*t = -30 sec*) and 30 min after (t = 30 min) wounding. Either in the wild type (WT, n = 9) PLCδPH-GFP or after knockdown of Rab35 (n = 9). Statistical significance was evaluated using the Wilcoxon matched-pairs signed rank-test. The following p-values were obtained: *p = 0.12* (ns), *p = 0.033* (*), *p = 0.002* (**) and *p < 0.001* (***). **(h)** Graphical summary of how Rab35 bridges actin and membrane dynamics in epidermal wound closure *in vivo* by regulating lamellipodia stabilization via Fascin and PIP2-delivery via vesicle trafficking (see also discussion). All experiments were performed from 3 independent biological replicates.

### Suppression of Rab35 function affects vesicular transport to the wound margin resulting in an accumulation of PIP2-positive vesicles

The localization of endogenous Rab35 in the wounded epidermis likely reflects a dual function of Rab35 in wound healing, as already suggested by our RNAi knockdown experiments. On the one hand, Rab35 can act through Fascin to promote the formation of cross-linked actin networks stabilizing lamellipodial protrusions. On the other hand, Rab35 could mediate the vesicular transport necessary for the effector proteins and expansion of the cell membrane at the wound site. To directly visualize membrane dynamics during wound closure we used the biosensor PLCδPH-GFP which selectively binds to the membrane phospholipid phosphatidylinositol 4,5-bisphosphate (PIP2). PIP2 is the most abundant phosphoinositide and is mainly localized in the plasma membrane but distinct functional pools also exist within the endosomal system^37^. It also functions as critical second messenger regulating diverse cellular activities, including reorganization of the actin cytoskeleton such as endosomal actin comet tail formation, endocytosis, exocytosis, and cell migration ^38^. In wild type LECs, the PLCδPH-GFP reporter localized in small puncta and along cell-cell interfaces on the apical surface, quite similar to endogenous Rab35 localization (Figure 3d; supplementary movie M10). Following an injury, the number of PLCδPH-GFP-labeled vesicular structures in cells at the wound margin decreased significantly, while at the same time there was a massive accumulation of the biosensor in newly formed lamellipodia (red arrowheads in Figure 3d, g; supplementary movie M10). This localization and dynamic distribution of the biosensor was significantly altered in cells depleted for Rab35 (Figure 3e, f; supplementary movies M11 and M12). Suppression of Rab35 function resulted in significant reduction of membrane enrichment of PIP2 and a marked increase in the number and size of PIP2-positive vesicles before wounding (Figure 3e, f; supplementary movies M11 and M12). After wounding, a large number of PIP2-positive vesicles were still detectable, although their number decreased as the wound partially closed. Overall, the strong accumulation of PLCδPH-GFP in protruding lamellipodia observed in wild type cells was significantly lower in Rab35 depleted cells.

These data suggest that Rab35 also controls the subcellular distribution of PIP2, an important second messenger and anchor of numerous signaling molecules and actin regulators that might contribute to efficient epidermal wound closure (Figure 3h).

## Discussion

Upon tissue damage, cells must coordinate actin-driven protrusions and membrane transport through localized mechanical/biochemical feedback and targeted vesicle trafficking. This long held hypothesis is based on numerous observations using diverse wound model systems ^5,8,24,39^. However, the molecular mechanisms and regulators that coordinate actin and membrane dynamics during epithelial wound healing are not well understood. To date, the role of some Rab proteins has been studied primarily in *in vitro* scratch-wound assays ^20,21^. In this study, we identified Rab35 as a novel regulator of the wound healing process with a dual capacity of regulating vesicle trafficking and modulation of actin dynamics. Rab35 acts through Fascin to stabilize actin networks in lamellipodial protrusions (Figure 3h). On the other hand, Rab35 controls the subcellular distribution of PIP2, an integral component of membranes, but also an important signaling molecule and activator of numerous actin regulators and cell adhesion molecules, including acting binding proteins including formin, gelsolin, cofilin, profilin, filamin, N-WASP, ezrin, α-actinin, and many others ^40,41^. PIP2 is the most abundant phosphoinositide and is mainly localized in the plasma membrane but also in endosomes ^37^. Suppression of Rab35 function resulted in significant reduction of membrane enrichment of PIP2. Instead, PIP2 was found to mislocalize into the intracellular vesicles. Interestingly, we also observed that F-actin often mislocalized on the large intracellular vesicles and numerous dynamic actin comet tails are found racing through epidermal cells depleted for Rab35. A likely hypothesis is that PIP2 progressively induces N-WASP-Arp2/3 mediated actin polymerization at the surface of these vesicles as shown in cultured cells ^42^. In these cell culture experiments, the type I phosphatidylinositol phosphate 5-kinase (PIP5KI) was overexpressed, which synthesizes PIP2, and promotes actin polymerization from membrane-bound vesicles to form motile actin comets. Future detailed studies are required to better understand how distinct phosphoinositide and lipid kinases differentially contribute to wound healing, both in a physiological and pathophysiological context ^43,44^.

An endosome-plasma membrane recycling activity was also the first function of Rab35 described in cytokinesis of *Drosophila* S2 cells ^45^. Similar to epithelial wound closure, cytokinesis depends on extensive localized membrane distortion and remodeling at the plasma membrane of the cleavage site that requires a polarized delivery of specific lipids and proteins. Thus, Rab35 might perform a recycling function analogous to that of Rab11 and its effectors ^46–48^. Notably, depletion of either Rab11 or Rab35 disrupts cytokinetic abscission and correlates with aberrant F-actin accumulation at the intercellular bridge ^49^. In our RNAi screen, we did not find Rab11 as a strong candidate but rather Rab2, Rab5, Rab7 and RabX4 (supplementary information) which will be described elsewhere.

The Rab35 activity is controlled by interaction with GEFs and GAPs ^50^. Here, we have identified the DENN-domain-containing protein CG18659 as a potential RAB35 GEF, the only ortholog of the three human DENND1 A-C genes in the fruit fly that binds to Rab35 and is also required for epidermal wound closure. Consequently, suppression of CG18659 and overexpression of Whacked, the orthologue of human TBC1D10A and known Rab35 GAP phenocopies depletion of Rab35 function. TBC1D10A and Rab35 regulate evoked exocytosis of endothelial Weibel–Palade bodies, a process that highly depends on intracellular Ca^2+^ elevation ^51^. One of the earliest conserved wound signals is the Ca^2+^ influx from the extracellular environment into damaged cells which might not only directly contribute to actin cytoskeleton but also membrane remodeling by Rab GTPases ^10,52,53^. Interestingly, Ca^2+^ was previously demonstrated to interact with the PIP2 headgroups and regulate the affinities of proteins to PIP2 ^54^. Moreover, some members of large Rab GTPases in mammals such as Rab44, Rab45/RASEF, and Rab46 contain an amino-terminal EF-hand domain that might act as novel membrane trafficking regulators with a calcium sensor function ^55^.

### Experimental procedures

#### Drosophila genetics

Flies were kept at room temperature in plastic vials, containing Drosophila standard food. For experimental procedures, fly husbandry and crossings were carried out according to standard methods and were kept at 25°C. In UAS-Gal4-based experiments, crossings were kept at 29°C. The following lines were obtained from Bloomington stock center (see also in supplementary information for all used RNAi fly stocks): UAS-LifeactGFP (BL57326), UAS-PLCδPH-GFP(BL39693), UAS-Rab35DN (BL9820), UAS-CG18659 RNAi (BL77436), UAS-wkd RNAi (BL77447) and w[1118] (BL3605). The A58-Gal4 driver was a kind gift from M. Leptin (Galko and Krasnow, 2004). UAS-Wkd was a kind gift from A. Ghabrial ^31^.

#### Plasmid construction

Rab35, a 606 bp PCR product, was amplified from cDNA LD21953 (DGRC) using the following primers: forward (5’–3’): cacc ATG GCA CGC GGC TTC GAT CAT and reverse (5’–3’): CCT GCA GCA CTT CCC ACC C. The Rab35 GEF CG18659, a 1824 bp PCR product, was isolated from cDNA LP09925 (DGRC) using the following primers: forward (5’–3’): cacc ATG GGA TCG CGT ATC AAA AAT G and reverse (5’– 3’): CTG GAG CTG CTG TAG CCG. PCR was performed using Q5 High-Fidelity DNA Polymerase (New England Biolabs). The PCR products were cloned into the entry vector pENTR-D-TOPO (Invitrogen). Orientation of the inserts was confirmed by sequencing using M13 forward and M13 reverse standard primers. Gateway cloning from pENTR-D-TOPO-Rab35 and pENTR-D-TOPO-CG18659 into pUASTattb-rfa-GFP or pUASPattb-rfa-myc was performed using LR-clonase mix (Invitrogen). Control digestions were done using EcoRV (New England Biolabs) for Rab35 or BamHI (New England Biolabs) for CG18659 followed by sequencing using the primers: pUAST-forward (5’–3’): GAA TAC AAG AAG AGA ACT CTG, pUAST-sv40-reverse (5’–3’): TAG AGC TTT AAA TCT CTG TAG G, pUASP-forward (5’–3’): CCT GTT CAT TGG TAC CCG, pUASp-kc10-reverse (5’–3’): GGT GCT ATG TTT ATG GCG CT.

#### Cell culture and transient transfection

Drosophila S2R+ cells (DGRC) were cultured in 1× Schneider’s medium supplemented with 10% fetal bovine serum, 50 U/mL penicillin, and 50 μg/mL streptomycin in 25 cm² cell culture flasks. For transient transfection, S2R+ cells were seeded into 24-well plates at a density of 5×10^5 cells per well. For each transfection, 0.6 μg Act5c-Gal4 and 1 μg plasmid DNA encoding the respective gene of interest under control of a UAS sequence were diluted in M3---medium and mixed with FuGENE®. The transfection mixture was incubated for 20 min at room temperature to allow formation of DNA-lipid complexes. Subsequently, the transfection mixture was added dropwise to each well. Transfected cells were incubated at 25 °C for 72 h.

#### Co-immunoprecipitation experiments and Western Blotting

For co-immunoprecipitation, Drosophila S2R+ cells (DGRC) were co-transfected with Rab35-GFP as well as CG18659-myc. The cells were lysed in ice cold lysis buffer containing 10 mM TrisHCl (pH 7.5), 150mM NaCl, 0.5 mM EDTA, 0.5 % NP40, 0.09 % NaN3. Lysates were incubated on ice, cleared by centrifugation and diluted in ice cold dilution buffer containing 10 mM TrisHCl (pH 7.5), 150 mM NaCl, 0.5 mM EDTA, 0.018 % NaN3 and protease inhibitors. GFP-tagged agarose beads (ChromoTek) were equilibrated in ice cold dilution buffer and sedimented by centrifugation. Diluted cell lysates were added to the equilibrated beads and incubated end-over-end for 2 hours at 4°C for protein binding. Afterwards, beads were washed with ice cold wash buffer containing 10 mM TrisHCl (pH7.5), 150 mM NaCl, 0.05 % NP40, 0.5 mM EDTA and 0.018 % NaN3. Bound proteins were eluted in Laemmli buffer and analyzed by Western blot. Equal amounts of protein (2% of cell lysates) were separated by SDS-PAGE and transferred to activated PVDF membranes (Thermo scientific). PVDF membranes were blocked in 10% milk in TBS-T and incubated with primary antibodies overnight at 4°C, followed by HRP-conjugated secondary antibodies. Detection was performed using chemiluminescence (ECL).

#### Immunofluorescence staining

Cells were plated on concanavalin A-coated coverslips, fixed with 4% paraformaldehyde, and shortly permeabilized with Triton X-100. Primary antibody incubation was performed, after blocking, for 2h, followed by incubation with fluorophore-conjugated secondary antibodies. F-actin was visualized using Alexa Fluor-488 or Alexa Fluor-568-conjugated Phalloidin (1:100 dilution, #A12379 #A12380, Invitrogen) and nucleus by DAPI staining (1 μg/mL, #62248, Thermo Scientific). Images were acquired using Leica confocal microscopes (SP8) and processed with ImageJ/FIJI. The following primary antibodies were used: anti-WAVE (affinity-purified guinea pig and rabbit (Bogdan et al., 2005); anti-Myc 9E10 (DSHB hybridoma supernatant); anti-GFP JL8 (Takara). Secondary antibodies conjugated to Alexa fluorophores were used according to standard protocols.

#### Quantitative analysis of wound closure, lamellipodia formation and PIP2-positive vesicles

Wound closure was evaluated by measuring the cell area using the polygonal selection tool in FIJI (ImageJ, NIH). Measurements were conducted at 5-minute intervals over a 30-minute period. The progress of wound closure was quantified as the relative reduction in wound area, determined by comparing the absolute wound area at each time point to the area measured at the time of maximum expansion (defined as 5 minutes post-wounding). The results were reported as a percentage of wound closure, with 0% representing the wound size at the time of maximum expansion and 100% representing complete closure. The lamellipodial area was assessed at 5-minute intervals over a 30-minute duration. The lamellipodial area was calculated by subtracting the area that was not covered by protrusion from the total wound area. In detail, a threshold-dependent binary mask was generated using FIJI (ImageJ, NIH) for each image. The area of the cell body was determined using the “Wand” tracing tool in FIJI (ImageJ, NIH), and the absolute lamellipodial area was subsequently derived by subtracting the cell body area from the total cell area. To quantify the lamellipodial response in relation to the injury, the lamellipodial area was expressed as a percentage of the total wound area. This was determined by normalizing the absolute lamellipodial area to the absolute wound area at each corresponding time point. The number of PIP2-positive vesicles at the apical epidermal surface was quantified via manual counting at two time points: before wounding and 30 minutes post-wounding. For each experimental recording, six cells were analyzed, consisting of the wounded cell and its five immediate neighbors.

#### Vesicle Tracking

To analyze vesicle dynamics, the “Manual Tracking” plugin in FIJI (ImageJ) was employed. A total of 30 vesicles per movie were tracked, and their trajectories were recorded. Based on the generated data, the direction of movement was classified manually as either toward or away from the wound edge.

#### In silico interaction modelling

To predict the binding interaction between Rab35 and CG18659, AlphaFold3 was employed. The respective amino acid sequences were retrieved from the UniProt database under the accession numbers Q7JXD2 and A1Z7Q5. Potential complex formation was modeled using default software parameters.

#### Statistics

All experiments were repeated in at least three independent (biological) replicates. Technical replicates were obtained from all experiments. Statistical analysis was performed with GraphPad Prism software Version 8. Wound healing experiments were analyzed using two-way ANOVA.

## Acknowledgements

We thank the Bloomington Stock Center and VDRC for fly stocks. We thank Katja Rust and Mila Y. Höhne for critical reading of the manuscript. The work was supported by grants to S.B. (BO1890/5-1 and BO1890/7-1) from the Deutsche Forschungsgemeinschaft (DFG).

## Data Availability Statement

All relevant data are in the manuscript and its Supporting information files.

## Author contributions

Sven Bogdan: Conceptualization, Project Administration, Funding Acquisition, Supervision, Methodology, Writing – Original Draft Preparation. Meike Schneider: Investigation, Methodology, Validation, Formal Analysis.

## Competing interests

The authors declare no competing interests.

## Supplementary movies

**Supplementary Movie 1**

Spinning disc microscope time-lapse movie of a wild type epidermis, expressing A58-Gal4, UAS-Lifeact-GFP (n = 9). F-actin accumulates at the wound margin, driving the formation of broad lamellipodial protrusions in adjacent apical epidermal cells. These protrusions reach their maximum extent approximately 10–15 minutes post-injury. Subsequently, lamellipodial activity subsides as contractile actin bundles assemble at the leading edge to facilitate the advancement of neighboring cells toward the wound site. Concurrently, an actomyosin cable develops at the basal side, acting as a ‘purse-string’ that draws opposing wound edges together. Furthermore, the accumulation of abundant vesicular structures strongly suggests that intracellular membrane trafficking plays an essential, parallel role in the wound healing response. Scale bar: 20 µm.

**Supplementary Movie 2**

Time-lapse movie from a spinning disc microscope showing single cell laser ablation in Rab35 RNAi (#1) under control of A58-Gal4 together with UAS-Lifeact-GFP. Rab35 knockdown (n = 9) leads to a marked reduction in cortical F-actin accumulation, the formation of actin-coated vesicular structures, and the emergence of highly dynamic actin comet tails within epidermal cells (yellow arrowheads). Compared to wild-type controls, r*ab35* depleted cells at the wound margin exhibit reduced lamellipodial protrusions and a significant delay in wound closure within 30 minutes post-injury. Scale bar: 20 µm.

**Supplementary Movie 3**

Live-imaging of single cell wounding in Rab35 RNAi (#2) under control of the A58-Gal4 driver combined with UAS-Lifeact-GFP. In *rab35* depleted cells a substantial decrease in cortical F-actin, alongside the emergence of numerous actin-coated vesicular structures and highly dynamic actin comet tails within the cytoplasm was observed. Adjacent cells at the wound margin displayed smaller lamellipodia and a significant delay in wound closure efficiency during 30 minutes post-injury compared to wild type controls. Scale bar represents 20 µm, n = 9.

**Supplementary Movie 4**

Time-lapse movie using spinning disc microscopy. An abdominal epidermis, overexpressing dominant-negative Rab35DN driven by A58-Gal4, UAS-Lifeact-GFP. Expression of Rab35DN (n = 9) resulted in a strong reduction in cortical F-actin, concurrent with the abnormal appearance of numerous F-actin-coated vesicles and highly dynamic actin comet tails within the cytoplasm of neighboring cells around the wound margin. Additionally, these surrounding cells at the wound margin exhibited cryptic lamellipodial protrusions and an impairment in wound closure efficiency during the initial 30 minutes post-wounding compared to the wild type control. Scale bar: 20 µm.

**Supplementary Movie 5**

Spinning disc microscope live-imaging movie of epidermal cell, expressing *fascin* RNAi under control of A58-Gal4 with UAS-Lifeact-GFP. Knockdown of *fascin* (n = 9) resulted in reduced lamellipodial protrusions and a significant prolongation of wound healing, compared to the wild type. Scale bar: 20 µm.

**Supplementary Movie 6**

Spinning disc microscope time-lapse movie of epidermal cells, expressing CG18659 RNAi under control of the A58-Gal4 driver in combination with UAS-Lifeact-GFP. Depletion of cg18659 recapitulated the cytoskeletal alterations observed following *rab35* knockdown, including the presence of actin comet tails and diminished lamellipodia. Scale bar represents 20 µm, n = 9.

**Supplementary Movie 7**

Live-imaging of single-cell laser ablation of an abdominal epidermis, expressing *whacked* RNAi under control of A58-Gal4, UAS-Lifeact-GFP. In contrast to *rab35* or *CG18659* depletion, knockdown of *whacked* (n = 9) did not affect wound healing. Cells around the wound margin exhibited large lamellipodia and displayed high wound closure efficiency compared to the wild type. Scale bar: 20 µm.

**Supplementary Movie 8**

Time-lapse movie of single-cell laser ablation using a spinning disc microscope. An abdominal epidermis, overexpressing Whacked, driven by A58-Gal4, UAS-Lifeact-GFP. In contrast to knockdown, overexpression of wild type Whacked (n = 9) resulted in reduced lamellipodial protrusions and a significant prolongation of wound healing, compared to the wild type. Scale bar: 20 µm.

**Supplementary Movie 9**

Time-lapse movie of single-cell laser ablation using airy scan confocal microscopy. An abdominal epidermis, expressing an endogenous Rab35-YFP protein. Before wounding, Rab35 exhibits a punctate distribution at cell-cell interfaces. Following tissue damage, Rab35 is recruited to vesicles accumulating at the wound site. Throughout wound closure, these vesicles fuse with the apical plasma membrane, while Rab35 also becomes increasingly concentrated within the protruding lamellipodia at the wound margin. Scale bar: 10 µm, n = 5.

**Supplementary Movie 10**

Spinning disc microscope time-lapse movie of a wild type epidermis, expressing A58-Gal4, UAS-PLCδPH-GFP. PLCδPH-GFP was used as a membrane-specific reporter to visualize PIP2. The reporter localizes in apical puncta and at lateral cell-cell contacts. Upon wounding, the number of PLCδPH-GFP-positive vesicles was decreased and PLCδPH-GFP assembled into broad lamellipodial protrusions. Scale bar: 20 µm, n = 9.

**Supplementary Movie 11**

Live-imaging of single-cell laser ablation of an abdominal epidermis expressing Rab35 RNAi (RNAi #1) under control of A58-Gal4, UAS-PLCδPH-GFP. Note: Suppression of Rab35 function resulted in significant reduction of membrane enrichment of PIP2 and a marked increase in the number and size of PIP2-positive vesicles even before wounding compared to wild type controls. Scale bar: 20 µm, n = 9.

**Supplementary Movie 12**

Live-imaging of single-cell laser ablation of an abdominal epidermis expressing a second *rab35* RNAi transgene (RNAi #2) under the control of the A58-Gal4 driver together with UAS-PLCδPH-GFP. Note: Suppression of Rab35 function resulted in significant reduction of membrane enrichment of PIP2 and a marked increase in the number and size of PIP2-positive vesicles even before wounding compared to wild type controls. Scale bar: 20 µm, n = 9.

## Supplementary information

**Supplementary table 1: Candidate genes selected for the RNAi-mediated knockdown screen.**

The *Drosophila* genome encodes 33 distinct Rab GTPases. To ensure phenotypic robustness and experimental reproducibility, 32 of these genes were targeted using two independent RNAi fly lines obtained from the Bloomington Drosophila Stock Center (BL) or the Vienna Drosophila Resource Center (VDRC). Specific genotypes and corresponding targets are provided. All UAS-transgenic lines were expressed under the control of the A58-Gal4 driver.

**Supplementary table 2: Results of the RNAi-mediated screen for Drosophila Rab GTPases**

The Drosophila genome encodes 33 distinct Rab GTPases. Individual RNAi-mediated knockdown resulted in a spectrum of phenotypes, which were categorized based on their severity. Candidates classified as exhibiting a ‘normal phenotype’ showed wound closure rates comparable to wild type controls (∼88 %). The ‘weak phenotype’ group was characterized by impaired wound closure (∼70 %) or reduced lamellipodial protrusion lengths. Finally, knockdown of Rab GTPases in the ‘strong phenotype’ group resulted in a pronounced decrease in wound closure (∼60 %) and significantly shortened lamellipodia.

## Notes

### Competing Interest Statement

The authors have declared no competing interest.

